# Dynamics of chromosome replication and its relationship to predatory attack lifestyles in *Bdellovibrio bacteriovorus*

**DOI:** 10.1101/519983

**Authors:** Łukasz Makowski, Damian Trojanowski, Rob Till, Carey Lambert, Rebecca Lowry, R. Elizabeth Sockett, Jolanta Zakrzewska-Czerwińska

## Abstract

*Bdellovibrio bacteriovorus* is a small Gram-negative, an obligate predatory bacterium that is largely found in wet, aerobic environments (i.e. soil). This bacterium attacks and invades other Gram-negative bacteria, including animal and plant pathogens. The intriguing life cycle of *B. bacteriovorus* consists of two phases: a free-living non-replicative attack phase wherein the predatory bacterium searches for its prey, and a reproductive phase, in which *B. bacteriovorus* degrades a host’s macromolecules and reuses them for its own growth and chromosome replication. Although the cell biology of this predatory bacterium has gained considerable interest in recent years, we know almost nothing about the dynamics of chromosome replication in *B. bacteriovorus.* Here, we performed a real-time investigation into the subcellular localization of the replisome(s) in single cells of *B. bacteriovorus.* Our results confirm that in *B. bacteriovorus* chromosome replication fires only during the reproductive phase, and show for the first time that this predatory bacterium exhibits a novel spatiotemporal arrangement of chromosome replication. The replication process starts at the invasive pole of the predatory bacterium inside the prey cell and proceeds until several copies of the chromosome have been completely synthesized. This chromosome replication is not coincident with the predator-cell division, and it terminates shortly before synchronous predator-filament septation occurs. In addition, we demonstrate that if this lifecycle fails in some cells of *B. bacteriovorus*, they can instead use two prey cells sequentially to complete their life cycle.

**Importance:** New strategies are needed to combat multidrug-resistant bacterial infections. Application of the predatory bacterium, *Bdellovibrio bacteriovorus*, which kills other bacteria including pathogens, is considered promising for bacterial infections. The *B. bacteriovorus* life cycle consists of two phases, a free-living, invasive attack phase and an intracellular reproductive phase, in which this predatory bacterium degrades the host’s macromolecules and reuses them for its own growth. To understand the use of *B. bacteriovorus* as a ‘living antibiotic’, it is first necessary to dissect its life cycle including chromosome replication. Here, we present for the first time a real-time investigation into subcellular localization of chromosome replication in a single cells of *B. bacteriovorus*. This process initiates at the invasion pole of *B. bacteriovorus* and proceeds until several copies of the chromosome have been completely synthesized. Interestingly, we demonstrate that some cells of *B. bacteriovorus* require two prey cells sequentially to complete their life cycle.

## Introduction

*Bdellovibrio bacteriovorus* is a small (0.2- to 0.5-μm wide and 0.5- to 2.5-μm long) Gram-negative bacterium that is unusual in its ability to invade and kill other Gram-negative bacteria. Moreover, it was demonstrated that *B. bacteriovorus* also benefits from interacting with Gram-positive biofilms (*Staphylococcus aureus*) (1). Bacteria belonging to the *Bdellovibrio* genus are largely found in wet, aerobic environments (i.e., soil) (2). *B. bacteriovorus* has received considerable research interest due to its intriguing life cycle and its great potential to be applied as an antimicrobial agent in industry, agriculture, and medicine. This bacterium proliferates within the periplasm of the prey cell and can invade a wide range of bacteria, including plant and animal pathogens (3–8).

*B. bacteriovorus* has a biphasic life cycle (see Fig. 6) that consists of: (i) a non-growing attack phase, in which a predatory bacterium finds a prey cell, attaches to its outer membrane and enters the periplasm; and (ii) a reproductive phase, in which *B. bacteriovorus* degrades the host’s macromolecules and reuses them for its own growth and chromosome replication. During the attack phase, *B. bacteriovorus* actively seeks the prey cell and is highly motile in liquid cultures due to the presence of a single polar, sheathed flagellum (9). The successful invasion of *B. bacteriovorus* requires that it adheres to the prey cell using its type IV pilus (10, 11), which is located at the pole opposite the flagellum. Thus, the predator cell has an asymmetry that resembles the polarity of *Caulobacter crescentus* cells (12). During the growth phase of *B. bacteriovorus*, the prey cell dies and is transformed into a spherical structure called a bdelloplast, and the predatory cell elongates inside the bdelloplast, forming a filament. At the end of the reproductive phase, this filament undergoes synchronous septation and progeny cells are released into the environment (13). Newly formed *B. bacteriovorus* escaped from bdelloplast go through a maturation phase where the cell length increases (13).

Although the cell biology of *B. bacteriovorus* has gained considerable recent interest (13–15) we know very little about the dynamics of chromosome replication in this predatory bacterium. *B. bacteriovorus* possesses a single circular chromosome (16) that contains all essential genes (e.g., those encoding the Dna proteins) and elements (e.g., an origin of chromosomal replication, *oriC*) required for its own replication. Genomic analysis revealed also the presence of SMC protein and ParAB*S* system (16), which are in other bacteria required for chromosomes segregation into daughter cells (17). RNA-seq analysis showed that the chromosome replication-related genes of *B. bacteriovorus* are actively transcribed during the reproductive phase and silenced during the non-growing, attack phase (18). Thus, the chromosome replication of *B. bacteriovorus* must be precisely coordinated with its unusual life cycle. It seems reasonable to assume that, as in other bacteria, the process is mainly regulated at the initiation step, which is a crucial cell cycle checkpoint. We recently characterized the key elements involved in the initiation of chromosome replication in *B. bacteriovorus* (19). We demonstrated that, as in other bacteria, *B. bacteriovorus* chromosome replication starts at a single origin of replication (named *oriC*). We showed that the replication initiator protein, DnaA, from *B. bacteriovorus* specifically binds and unwinds its own *oriC in vitro* and *in vivo* (19). Beyond this, however, regulation of replication and the dynamics of this process during the *B. bacteriovorus* cell cycle are still unknown.

Recent studies have shown that the positioning of replisomes (i.e., the replication machinery) and their dynamics during the cell cycle differ among bacteria. In some bacteria (*Bacillus subtilis, E. coli* and *P. aeruginosa),* the replisomes are assembled in the middle of the cell whereas in others (C. *crescentus, H. pylori,* and chromosome I of *Vibrio cholerae*) this assembly occurs at one of the cell poles. During the replication cycle, the sister replisomes may stay together at the initiation site (*B. subtilis* and *P. aeruginosa*) or travel together to the midcell (C. *crescentus* and *H. pylori*) (20–27). While in *E. coli,* the sister replisomes move toward the cell poles and merge again at the end of replication (24). Recent work has shown that replisome dynamics may exhibit other patterns, such as those seen for *Mycobacterium smegmatis* and *Myxococcus xanthus* (28, 29), suggesting that bacteria evolve different replication fork passage strategies that are coupled to their specific life cycle requirements.

In this study, we addressed for the first time how the dynamics of chromosome replication are coordinated with the life cycle of *B. bacteriovorus*. We investigated the subcellular localization of the replisome(s) in real time in single cells/filaments of *B. bacteriovorus*. Our data provide evidence that *B. bacteriovorus* exhibits a novel spatial arrangement of chromosome replication. The process starts at the invasive pole of the predatory bacterium, inside the bdelloplast, and replication proceeds until several copies of the entire chromosome are completely synthesized. This chromosome replication is not associated with cell division, and it is terminated before synchronous predator-filament septation. In addition, we observed (albeit rarely) that some *B. bacteriovorus* cells do not follow a canonical life cycle, but rather employ two prey cell invasions to complete their life cycle if the first predation event is abortive. We also discover that some predatory *Bdellovibrio* cells are still competent to invade prey when their replisome is pre-assembled.

## Results

### Replisomes are formed during the reproductive phase

In recent years, the application of fluorescence microscopy has allowed direct observation of the replication dynamics in single bacterial cells in real time. The replisome is usually visualized by using fluorescent proteins fused to various subunits of the DNA polymerase III holoenzyme, including the sliding clamp (β-subunit) protein, DnaN (30–33, 29). To monitor the positioning of the replisome in *B. bacteriovorus* cells, we constructed strain HD100 DnaN-mNeonGreen/PilZ-mCherry, which produced the two fusion proteins, DnaN-mNeonGreen and PilZ-mCherry (Fig. S1 and see Materials and Methods). PilZ (Bd0064; a protein that binds cyclic di-GMP) is localized nearly constitutively (34) throughout the cytoplasm of the *B. bacteriovorus* cell, so its fluorescent tagging allowed us to label the entire predatory cell in red (Fig. 1I and Fig. S1B). Ongoing replication was visualized by the appearance of DnaN-mNeonGreen foci. These signals enabled us to precisely monitor the position of replisome(s) inside a predatory cell growing in the bdelloplast. The DnaN-mNeonGreen/PilZ-mCherry strain exhibited a predatory kill curve and predation efficiency similar to those of the wild-type strain (Fig. S2), suggesting that the fusion proteins were fully functional.

**Figure 1.**
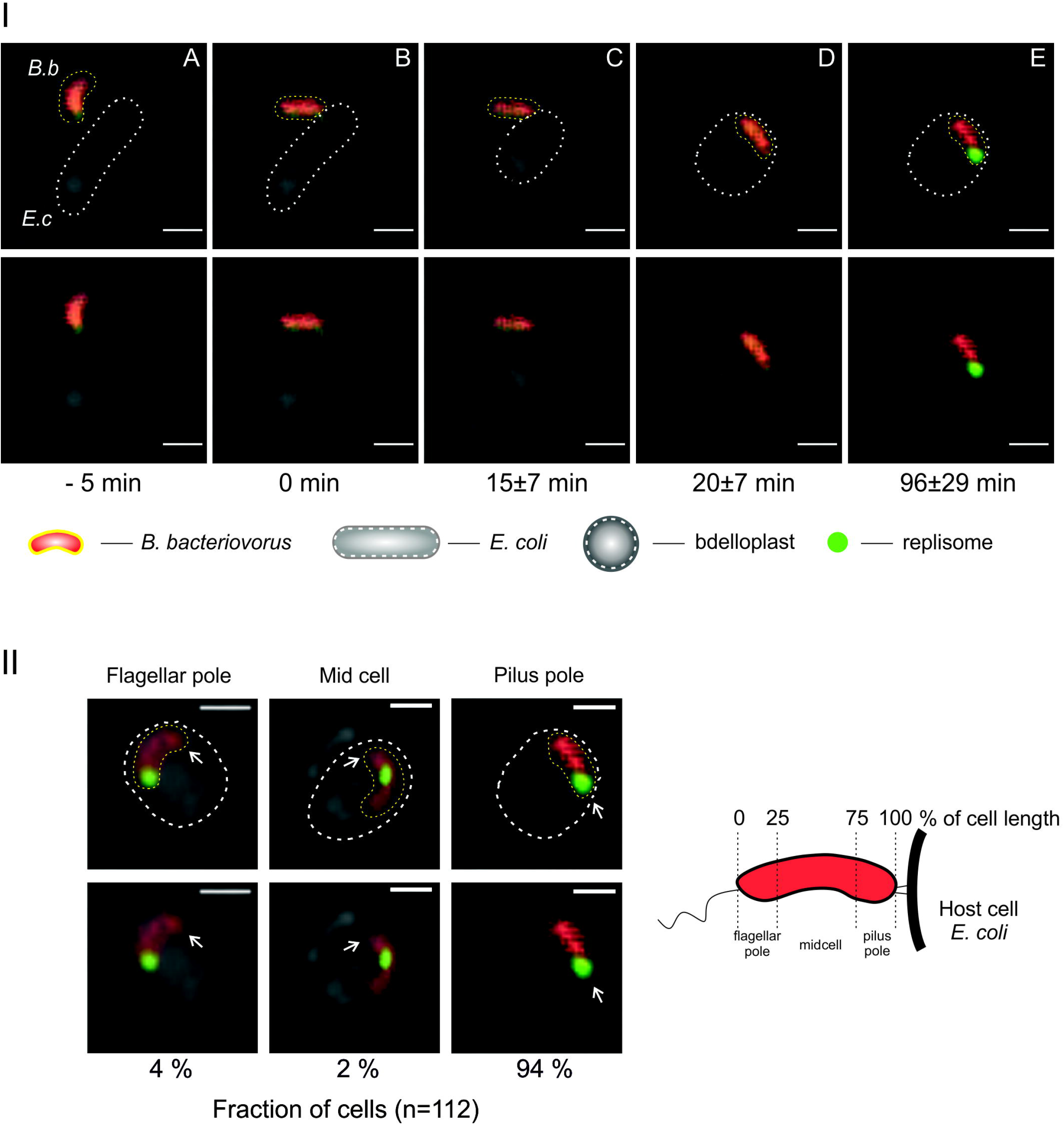
Spatiotemporal analysis of replication initiation in *B. bacteriovorus*. (**I**) Time-lapse analysis of the appearance of the first replisome (green) in *a B. bacteriovorus (B.b)* cell (red) growing inside an *E. coli (E.c)* bdelloplast (**A-E**). Predatory attachment to the host cell corresponds to time = 0 min. (**II**) Localization of the first replisome (green) in relation to the pilus pole of *B. bacteriovorus* (red). Cells with replisome localized in the vicinity of a pole (flagellar or pilus) or in the midcell are shown. The white arrow indicates the pilus pole, determined by watching predator entry, which is pilus first. A schematic of a *B. bacteriovorus* cell is depicted on the right. The cell is divided into subregions according to the percentage of cell length. Images were recorded every 60 seconds. Red - PilZ-mCherry labelled cytoplasm of *Bdellovibrio* and green - DnaN-mNeonGreen of *Bdellovibrio.* All photos represent merged DIC (differential interference contrast) and fluorescence (red and green) images. The *B. bacteriovorus* and *E. coli* cells are marked by yellow and white dotted lines, respectively. Scale bar = 1 μm.

To analyze the duration and timing of *B. bacteriovorus* chromosome replication, we used an agarose pad in combination with ibidi cell-imaging dishes (see Materials and Methods). In this system, the predatory cells could move around beneath the agarose pad, whereas the immobilized prey cells (i.e., *E. coli*) were able to form bdelloplasts. We were thus able to observe the complete life cycle of *B. bacteriovorus*.

Preliminary microscopic analysis revealed that DnaN-mNeonGreen fluorescence was visible either as a dispersed signal found throughout the cell during the (non replicating) attack phase (Fig. S1B) or as discrete diffraction-limited foci observed during the reproductive phase (inside the bdelloplast) (Fig. 1 IE). In the latter case, we observed up to four DnaN-mNeonGreen foci per single *B. bacteriovorus* filament (Fig. 2). The presence of diffuse DnaN-mNeonGreen fluorescence during the attack phase likely reflects that the replication machinery is disassembled in newborn cells.

**Figure 2.**
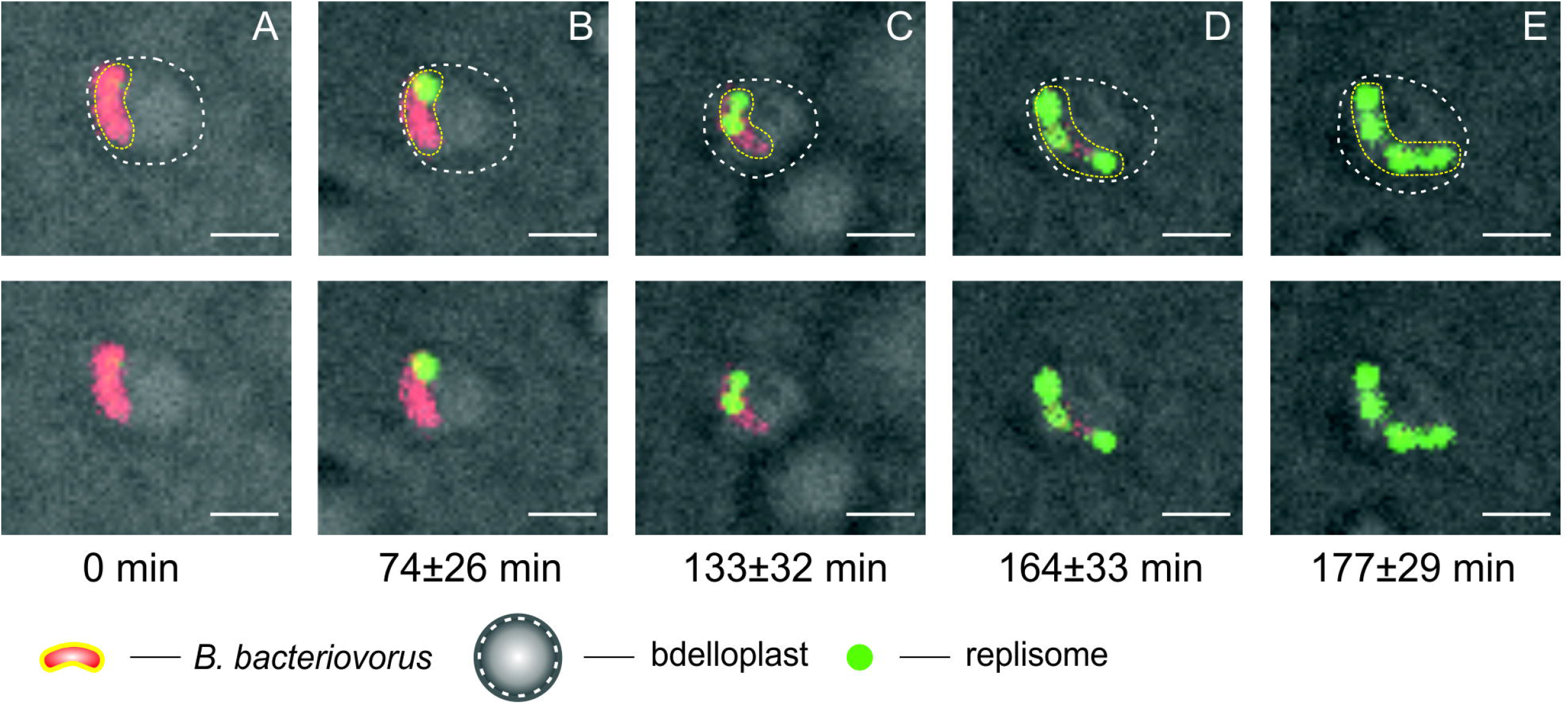
Dynamics of chromosome replication in a *B. bacteriovorus* cell growing in a bdelloplast. Time-lapse analysis showing the localization of replisomes (green) in a *B. bacteriovorus* cell (red) growing inside the *E. coli* bdelloplast (**A-E**). Time = 0 min indicates the initial formation of the bdelloplast. Red - PilZ-mCherry labelled cytoplasm of *Bdellovibrio* and green - DnaN-mNeonGreen of *Bdellovibrio.* Photos represent merged DIC and fluorescence (red and green) images. The *B. bacteriovorus* cell and the bdelloplast are marked by yellow and white dotted lines, respectively. Scale bar = 1 μm.

To examine whether the observed fluorescent foci legitimately reflected ongoing replication, we used novobiocin; this agent inhibits DNA replication by acting on DNA gyrase, which normally, through relaxation of positive supercoils ahead of the replication fork, resolves the torsional tension and allows DNA synthesis to be continued (35). Indeed, when *B. bacteriovorus* cells were treated with novobiocin present in 1% agarose pad (200 μg/ml), the fluorescent foci disappeared and only diffuse fluorescence was observed (Fig. S1C). This confirms that the DnaN-mNeonGreen foci represented active replisomes.

### Chromosome replication starts at the invasive pole of the *B. bacteriovorus* cell and two or more replisomes are usually observed in a single filament

Time-lapse analysis of the DnaN-mNeonGreen/PilZ-mCherry strain of *B. bacteriovorus* revealed that most of the cells growing inside the bdelloplast contained two (28%), three (46%) or four (25%) visible replisomes (Table 1). Only a small fraction of cells (1%) contained more than four replisomes. As would be expected, longer filaments of *B. bacteriovorus* usually contained more replisomes than shorter ones. Replisomes position (except for that of the first replisome, see below) were not restricted to specific cell regions (Fig. 2E). During the late stage of *B. bacteriovorus* cell growth, the filament can reach a length exceeding the bdelloplast diameter and begin to curve and overlap itself (13, 15, 36). In such cases, it becomes difficult to analyze the filament length and replisome number.

**Table 1.** Number of visible replisomes in growing *B. bacteriovorus* cell

| No. of visible<br>replisomes | No. of cells | Fraction [%] |
| --- | --- | --- |
| 2 | 31 | 28% |
| 3 | 51 | 46% |
| 4 | 28 | 25% |
| > 4 | 1 | 1 % |

A *B. bacteriovorus* cell enters a prey cell by using the type IV pili located on the non-flagellate pole of the predatory bacterium (11, 37). Careful tracking of predatory entry into *E. coli* cells allowed us to observe the appearance of the first focus (i.e., replisome) in relation to given cell pole of *B. bacteriovorus.* The predatory cells were acquired every 60 seconds using time-lapse fluorescence microscopy (TLFM). The TFLM analysis showed that *B. bacteriovorus* cells after entering *E. coli* did not flip inside the prey’s periplasm (n=90, see also Fig. S3 and Movie S1). Microscopic analysis revealed that in 94% (n=112, Fig. 1 II) of cells, the first replisome was assembled at the invasive (pilus-proximal) pole of the cell. In a small fraction of cells (6%), the first replisome was observed either at the flagellar pole or at the midcell (Fig. 1 II). TLFM analysis revealed that the first fluorescent focus appeared at 96±29 min (n=112) after the attachment of *B. bacteriovorus* to the *E. coli* cell (Fig. 1 IE) and 74±26 min (n=112) after bdelloplast formation (Fig. 2B). The time intervals between the appearances of consecutive replisomes varied (Table S2): the second fluorescent focus was assembled 59±20 min (n=111) after appearance of the first, while the third and fourth replisomes appeared (when relevant) after shorter time intervals of 32±18 min (n=80) and 27±15 min (n=28), respectively (Table S2). Although the *B. bacteriovorus* filament could be visualized inside the bdelloplast, it was difficult to determine the positions of replisomes within a growing *B. bacteriovorus* filament because they were highly mobile and frequently mixed with each other (Movie S2).

In summary, our results indicate that *B. bacteriovorus* chromosome replication is initiated at the formerly piliated invasion predator pole.

### The number of progeny cells is proportional to the duration of chromosome replication

To determine the duration of DNA replication (the C period) during the growth phase of *B. bacteriovorus*, we measured the time from the appearance of the first focus/replisome (regarded as initiation) to the disappearance of the last focus/replisome (estimated to be termination) (Fig. 3 I). The average duration of chromosome replication was 144±26 min (range 112 to 187 min, n= 112; Fig. 3 IC and Table 2). Because the length of the C period varied significantly between bdelloplasts, we analyzed the relationship between the number of progeny cells released from the bdelloplasts and the duration of DNA replication (*B. bacteriovorus* growing in abnormally elongated host cells, see below, were excluded from the regression analysis). As expected, the number of progeny cells was positively correlated with the duration of *B. bacteriovorus* chromosome replication (correlation coefficient, R^2^=0.97; Fig. 3II). In all cells, the replication process was terminated 21 min (or shorter, n=98), prior filament septation (D period).

**Figure 3.**
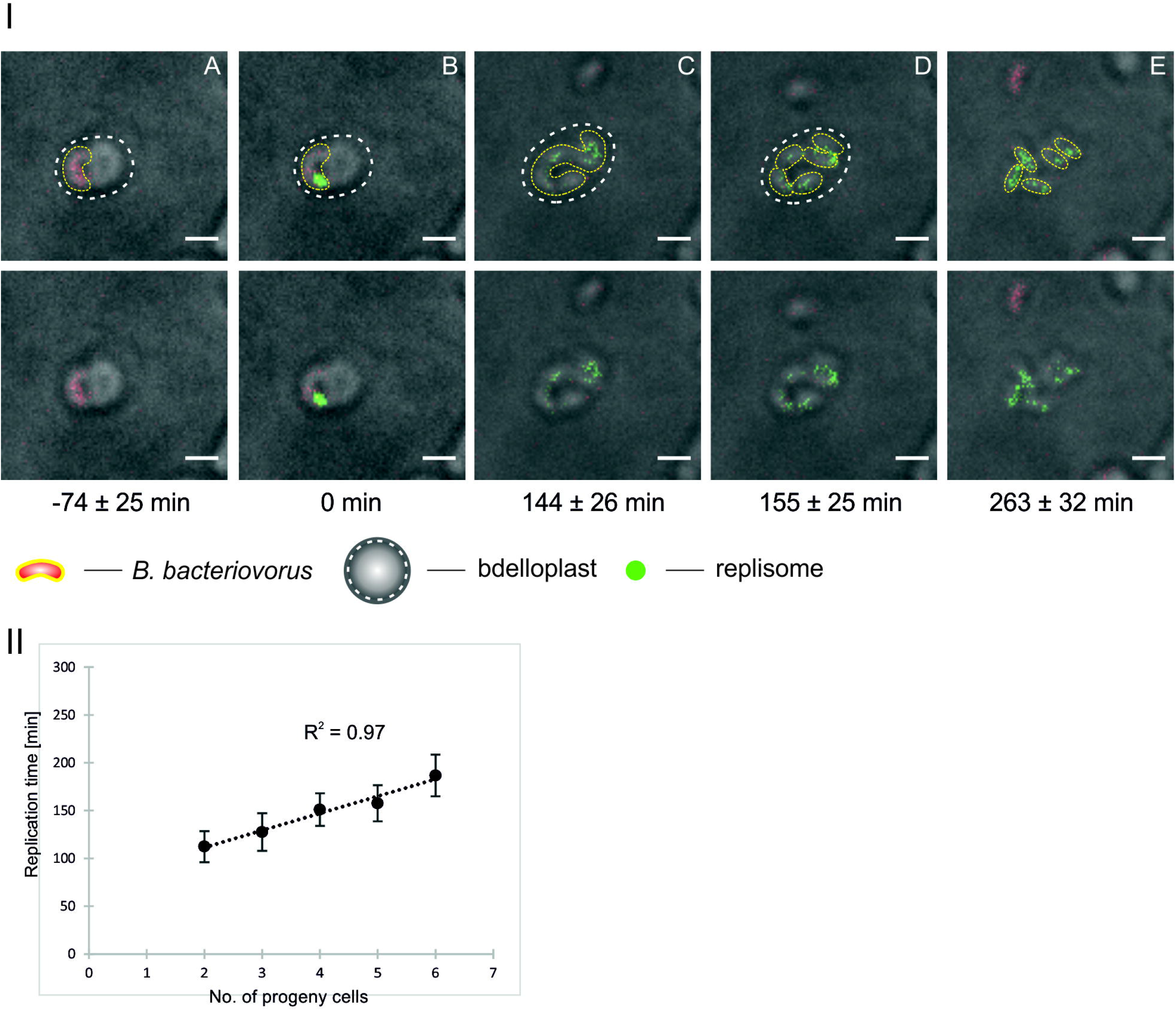
Duration of chromosome replication and its correlation to the number of *B. bacteriovorus* progeny cells. (**I**) Time-lapse analysis of the duration of *B. bacteriovorus* chromosome replication. Bdelloplast formation (**A**). The start of chromosome replication, time = 0 min (**B**). Replication termination (**C**). The beginning of *B. bacteriovorus* filament septation (**D**) and the release of progeny cells from the bdelloplast (**E**). Red - PilZ-mCherry labelled cytoplasm of *Bdellovibrio* and green - DnaN-mNeonGreen of *Bdellovibrio.* Photos represent merged DIC and fluorescence (red and green) images. The *B. bacteriovorus* cell and the bdelloplast are marked by yellow and white dotted lines, respectively as determined by careful analysis of DIC images. Scale bar =1 μm. (**II**) Graph presenting the correlation between the number of progeny cells and the duration of chromosome replication. Correlation coefficient, R^2^ = 0.97, n= 112.

**Table 2.** Duration of chromosome replication in different progeny cells

| No. of<br>progeny<br>cells | Free-living cells |  | Newly-released cells |  |
| --- | --- | --- | --- | --- |
|  | Duration of<br>replication<br>[min] | Standard<br>deviation | Duration of<br>replication<br>[min] | Standard<br>deviation |
| 2 | 112 | 16 | - | - |
| 3 | 128 | 20 | 122 | 11 |
| 4 | 151 | 17 | 140 | 14 |
| 5 | 158 | 19 | 147 | 23 |
| 6 | 187 | 22 | 170 | 7 |

During our TLFM analyses, we observed that *E. coli* occasionally formed extremely elongated cells (Fig. S4A) and that *B. bacteriovorus* easily invaded these cells and formed a huge, oval bdelloplast (Fig. S4B). As noted by Kessel and Shilo (36), we observed that the *B. bacteriovorus* filament reached an abnormal length in such bdelloplasts as there will be a greater nutrient supply. The elongated filament contained numerous replisomes (up to six, see Fig. S4E) that were evenly positioned within the growing predatory filament and appeared sequentially (Movie S3). In one particular case, replication lasted longer (285 min) and surprisingly ended simultaneous to septation (Movie S3). Twelve progeny cells were released from this abnormal bdelloplast (Fig. S4F and Movie S3).

### Replication begins earlier in progeny predator-cells that immediately invade new prey cells than in free-living predatory cells that invade prey cells

After being released from the bdelloplast, a progeny cell might attack prey in the surrounding neighborhood or actively move to more distantly located prey, taking several minutes to do so. We observed that if a newly-released progeny cells were in the close vicinity of another prey cell, they could quickly attack these preys (within 10±5 min; n=39) and form a new bdelloplast. In such bdelloplasts, replication started significantly earlier (23±11 min) than in the case of bdelloplasts formed by free-living mature predatory cells (74±26 min) (p-value<0.001, n=39; Table S2), but we did not noticed any significant differences in the duration of replication between newly-released predatory cells (140±20 min) and free-living cells (144±26 min)(p>0.05). Moreover, as in mature, free-living *B. bacteriovorus* cells (Table 2), in newly-released cells, the length of the C period was positively correlated with the number of progeny cells (data not shown). Surprisingly, in a few newly-released predatory cells (10%, n=39), the replisome was still visible after septation and bdelloplast lysis (Fig. 4B). In addition, one of these cells invaded the next prey and even during the invasion, the replisome was still observed (Fig. 4C and Movie S4). In this cell, the second replisome appeared already 35 min after the bdelloplast was formed (Fig. 4E) indicating that the DNA replication had to begin earlier.

**Figure 4.**
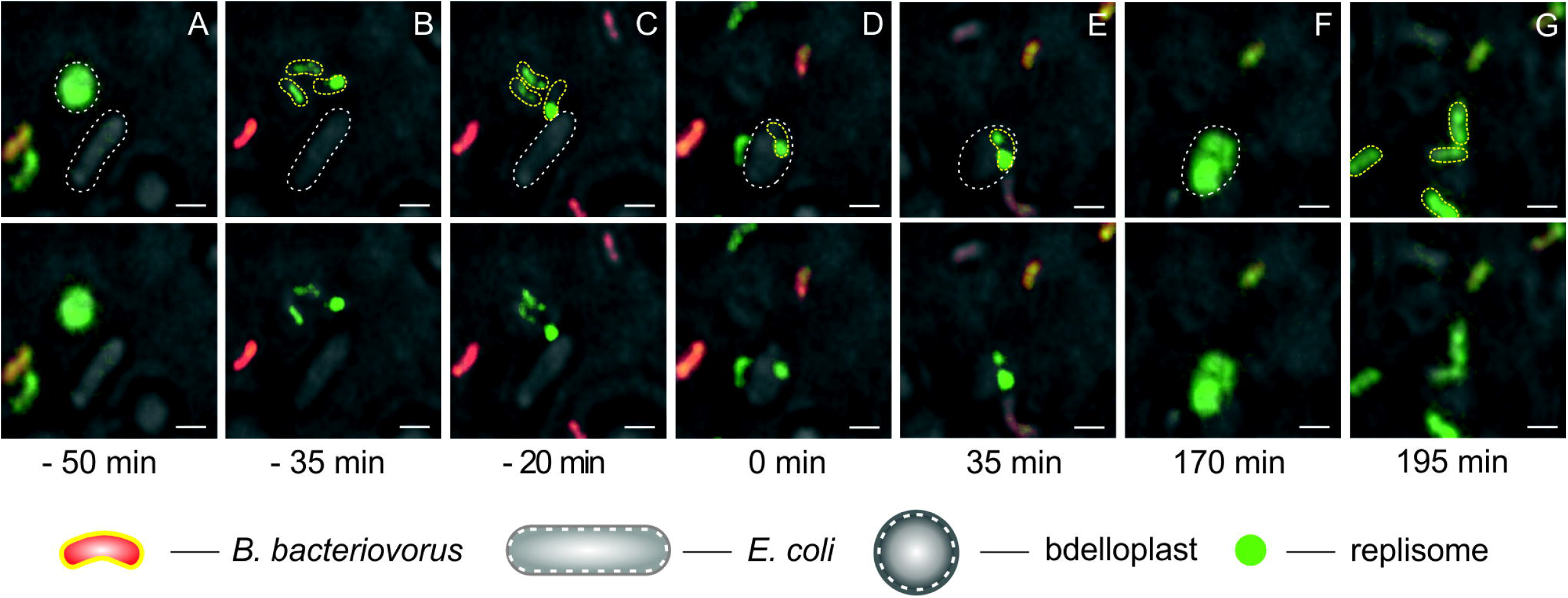
Prey predation by *B. bacteriovorus* cell with visible replisome at cell pole in the attack phase. Prior bdelloplast septation (**A**). Prior bdelloplast lysis releases new progeny cells (**B**). Attachment of a newly-released *B. bacteriovorus* - still with visibly-labelled replisome focus at the pilus pole to an *E. coli* cell (**C**). Rounding of this invaded *E. coli* cell to form bdelloplast, t = 0 min (**D**). Appearance of second replisome focus in predator indicates rapid restart of replication (**E**). Termination of predator chromosome replication (**F**). Bdelloplast lysis and release of progeny cells (**G**). Red - PilZ-mCherry labelled cytoplasm of *Bdellovibrio* and green - DnaN-mNeonGreen of *Bdellovibrio.* Photos present merged DIC and fluorescence (red and green) images. The *B. bacteriovorus* and *E. coli* cells are marked by yellow and white dotted lines, respectively. Scale bar = 1 μm.

These findings indicate that newly-released progeny cells that rapidly invade new prey cells show an earlier initiation of chromosome replication compared to free-living *B. bacteriovorus* cells that invade prey cells suggesting a time course of resetting to nonreplicative attack phase after prey exit.

### Some *B. bacteriovorus* cells might require two prey cells to complete their life cycle

Careful TLFM analysis of *B. bacteriovorus* cells allowed us to observe predator cell that did not complete its life cycle within the *E. coli* bdelloplast. Consistent with results published by Fenton *et al.* (13), we noticed that the filament failed to undergo septation. The undivided filament exited the bdelloplast (whether actively or passively is the subject of further research beyond this paper) and encountered another prey within which it completed its life cycle (Fig. 5). In such “two-stage” growing phase, the predator cell replicated its chromosome inside the first bdelloplast but retained visible replisomes for a much longer period than seen in the normal life cycle (390 vs. 187 min; Fig. 5D). The replisome was observed in the filament even after its release from the first prey cell (Fig. 5E and Movie S5). Upon its release, this elongated *B. bacteriovorus* filament entered new prey cell, whereupon chromosome replication proceeded (lasting 365 min) (Fig. 5H-I and Movie S5). Growth in a second prey cell ended with filament septation (Fig. 5J).

**Figure 5.**
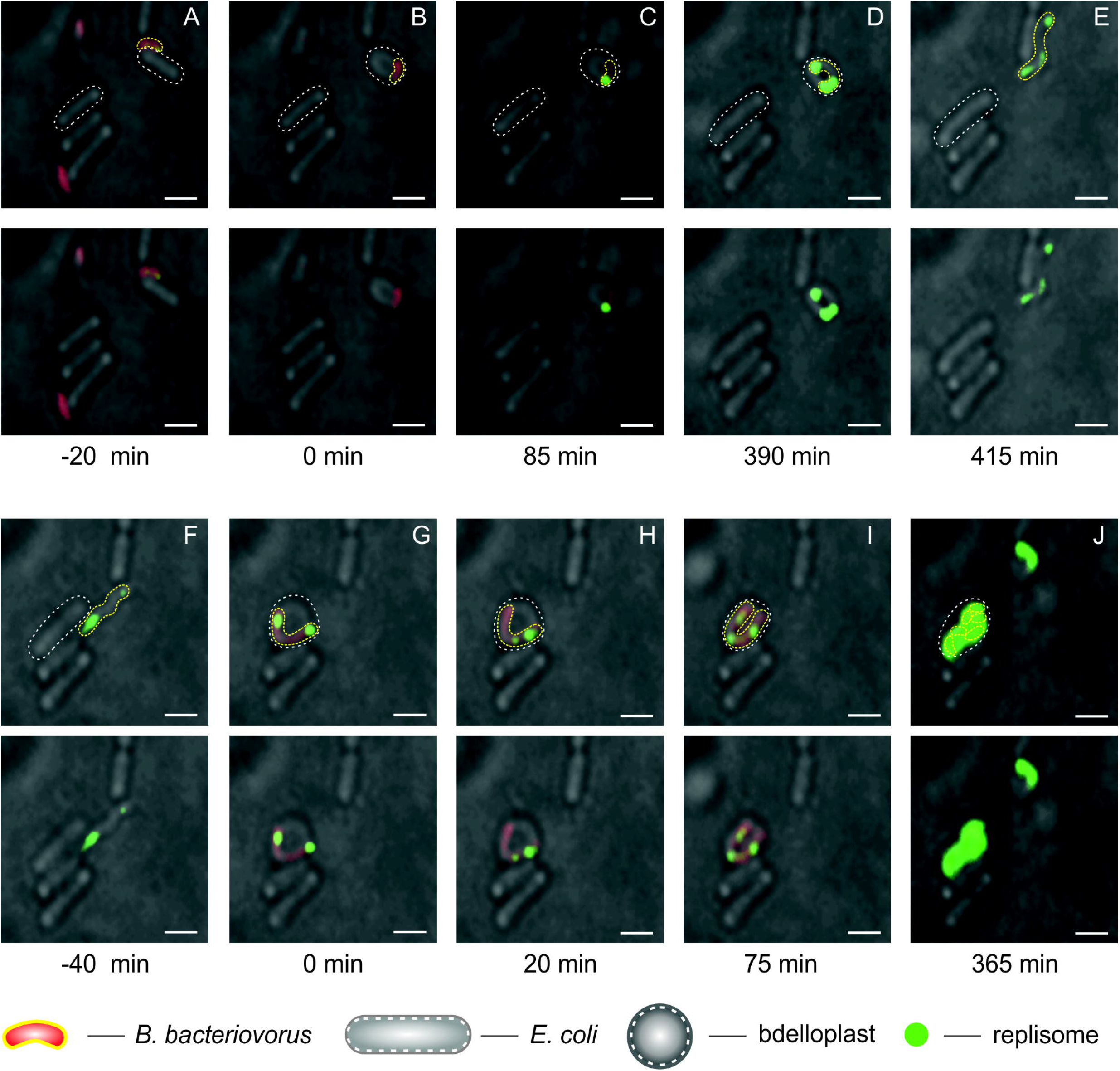
A rare example of a *B. bacteriovorus* cell life cycle conducted in two independent *E. coli* host cells. *B. bacteriovorus* attachment to *E. coli* (**A**). Bdelloplast formation, time = 0 min (**B**). Growth and replication in the first host cell (**C-D**). Novel release of the non-septated predatory filament from the first host cell (**E**). The attack of a non-septated predatory filament on another prey cell (**F**). Bdelloplast formation, t = 0 min (**G**). Growth and replication in the second host (**H** and **I**); Filament septation inside the bdelloplast (**J**). Red - PilZ-mCherry labelled cytoplasm of *Bdellovibrio* and green - DnaN-mNeonGreen of *Bdellovibrio.* Photos represent merged DIC and fluorescence (red and green) images. The *B. bacteriovorus* and *E. coli* cells are marked by yellow and white dotted lines, respectively. Scale bar = 1 μm.

These observations indicate that rarely *B. bacteriovorus* cell might require two independent prey cells to complete their life cycle.

## Discussion

The chromosome replication of *B. bacteriovorus* occurs only during the reproductive phase within the prey while the motile, free-living cells are incapable of initiating of chromosome replication. Thus, as in *C. crescentus,* the chromosome replication process must be strictly regulated and coordinated with the unusual life cycle of this predatory bacterium (3, 36) (see Fig. 6). Although chromosome replication dynamics have been relatively well studied in several species of Gram-positive and Gram-negative bacteria, almost nothing is known about this process in predatory bacteria. To address this, we developed a TLFM-based system that allowed us to observe chromosome replication dynamics in a single cell of *B. bacteriovorus* growing inside the prey bacterium, *E. coli.* Here, we report that this predatory bacterium exhibits a novel spatiotemporal arrangement of chromosome replication dynamics. Moreover, we found that *B. bacteriovorus* cells can use two independent prey cells to complete its life cycle if the first predation event fails.

**Figure 6.**
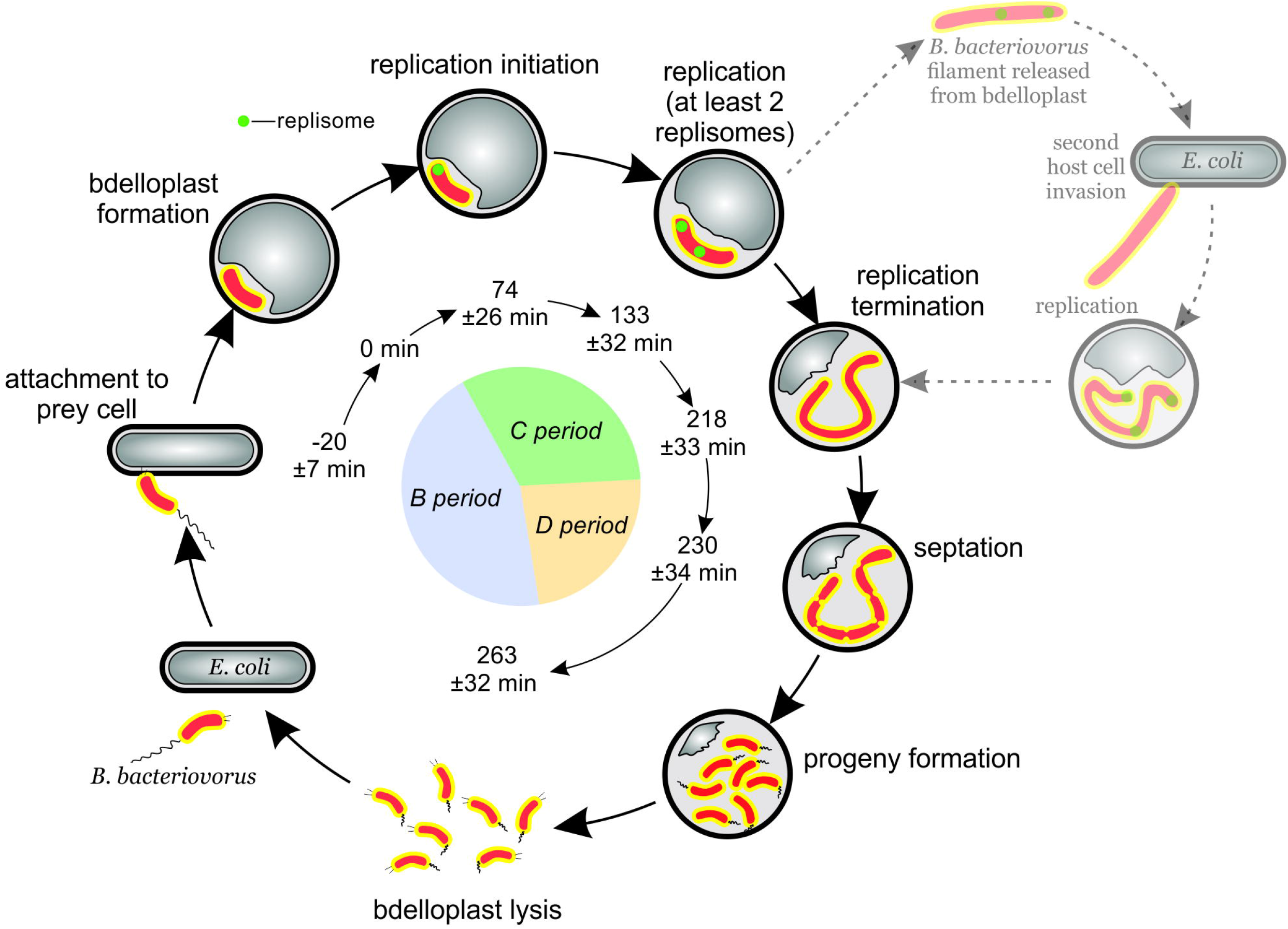
Dynamics of chromosome replication during the *B. bacteriovorus* life cycle. *B. bacteriovorus* (orange) attacks and invades the host cell (gray). Chromosome replication (green replisome) is initiated at the invasive pilus-proximal pole. The dotted line and greyscale represent the speculated alternative way of *B. bacteriovorus* life cycle conducted in two independent host cells. The inner circle diagram represents the period of the bacterial cell cycle (B, C, D); t = 0 min refers to bdelloplast formation. The listed time points were calculated in this study.

Our data indicate that the chromosome replication of *B. bacteriovorus* starts at the invasive pole (Fig. 1II). This pole is essential for predation, especially the entry of this bacterium into its prey (11). Moreover, a regulatory protein hub controlling predatory invasion was discovered at this pole (38). The pili protruding from the invasive pole take part in sequentially sensing the stepwise phase transition in *B. bacteriovorus* (38, 39). During prey recognition, the pili mediate the transduction of a yet-unidentified early signal that occurs in the cytoplasmic membrane of the host (38). The second cue, which also has not yet been specified, originates from the prey cytoplasm and is believed to promote DNA replication (39). Thus, the invasive pole of *B. bacteriovorus* seems to be involved in the transition from the attack phase to the reproductive phase. We speculate that during the transition phase, chromosome replication is triggered by yet-unknown regulator(s), presumably by signal transduction cascade(s), and that this process is likely to be mediated by the recognition of the cue arising from the invasive pole (38, 39).

Unlike *B. bacteriovorus*, model bacteria such as *E. coli* and *B. subtilis* undergo replisome assembly in the middle of the cell (22, 24, 25). Interestingly, *C. crescentus* and *V. cholerae* (chromosome I) resemble *B. bacteriovorus* both in their asymmetry and in assembling their replisomes at a cell pole (21, 40). In *C. crescentus* and *V. cholerae* (chromosome I), the subcellular localization of *oriC* (and thus the sites of replisome assembly) is determined by the specific oriC-anchoring proteins, PopZ and HubP, respectively (41, 42). The factor(s) responsible for anchoring the *B. bacteriovorus oriC* region at the invasive pole remains to be identified.

Spatiotemporal analysis of the chromosome dynamics in *B. bacteriovorus* revealed that the first replisome appears 96±29 min and 74±26 min after the attachment of the predatory cell to the prey cell (Fig. 1I) and the formation of the bdelloplast (Fig. 2), respectively. This pronounced delay in the initiation of chromosome replication (replisome assembly) presumably reflects the unusual predatory behavior of *B. bacteriovorus*. After entering a prey cell, the predatory cell must adapt to growing in the bdelloplast before it can begin DNA replication. Indeed, meta-analyses of gene expression profiles (RNA-seq and microarray profiling) have demonstrated that during the first 60 min post infection, genes involved in growth and replication are highly upregulated (43, 18). Our present results show that DNA is not yet being synthesized at this time (Fig. 1). Thus, we speculate that as yet undiscovered replication checkpoints act to coordinate the cell cycle progression and DNA replication of *B. bacteriovorus.* The predatory cell modifies the structure of the host’s peptidoglycans to make the environment more flexible and suitable for filamentous growth (44–49). Moreover, *B. bacteriovorus* during the adaptation inside bdelloplast releases hydrolytic enzymes to the prey’s cytoplasm to degrade various prey macromolecules and uses these components to build its own cellular structures (18). The chromosome replication of *B. bacteriovorus* is assumed to be triggered only after the bacterium adapts to the growth conditions inside the bdelloplast.

Using bdelloplasts that produced only two progenitor cells (11%; n = 112), we were able to calculate the rate of DNA synthesis. Given the length of the C period for such cells (112 min), the rate of DNA synthesis is about 300 nt/s. This is ~2–3 times slower than that of *E. coli* (600 – 1 000 nt/s; (50)). The activity of *B. bacteriovorus* DNA polymerase III is not likely to be the rate-limiting factor, since subunits of the holoenzyme show high homology (Fig. S5 and Fig. S6) with the corresponding subunits from *E. coli* (crucial amino acids for catalytic activity of α subunit of DNA polymerase III are identical in *B. bacteriovorus* and *E. coli*; see Fig. S6). Thus, the DNA synthesis rate of *B. bacteriovorus* is presumably limited by the availability of nutrients, particularly nucleotides (see below).

In filaments that formed more than two progeny cells, the C period ranged up to 187 min, indicating that in these filaments, reinitiation of chromosome replication must take place; to synthesize three or more chromosomes within less than 187 min, a new round of replication must be initiated before the previous round is completed. Thus, the reinitiation mechanism ensures that each of the nascent progeny cells receives a single chromosome.

The duration of the growth phase including the C-period in *B. bacteriovorus* varies between cells, but it is not yet known how the length of this phase is regulated. Gray and Ruby (51) suggested that prey-derived regulatory factor(s) may be involved in the developmental cycle of *B. bacteriovorus,* operating at the level of the cell’s decision to either continue or terminate the growth phase. As in other bacteria, *B. bacteriovorus* presumably adjusts its size and growth rate according to the availability of nutrients. Indeed, a *B. bacteriovorus* cell that attacks a large (i.e., large nutrient pool) prey cell will synthesize more chromosomes and develop a longer filament ((36) and Fig. S4) and thus release more progeny cells (Fig. S4F). The predatory cell to synthesize 2-3 nascent chromosomes utilizes DNA and RNA of prey as are direct sources of nucleotides (52, 53), but synthesis of more chromosomes (and consequently progeny cells) requires *de novo* synthesis of nucleotides from carbon and nitrogen precursors including amino acids obtained by hydrolysis of prey’s proteins (53).

Surprisingly, we observed some *B. bacteriovorus* cells in which replication was initiated relatively shortly after their invasion into new prey cells. This occurred only among newborn predatory cells that were released in close proximity to new prey cells and invaded them immediately upon release. In such predatory cells, replication began significantly earlier than in free-living predatory cells that underwent invasion (23 min versus 74 min, respectively, p<0.001; Table S2). It can be assumed that the proteins involved in chromosome replication (e.g., the initiator protein, DnaA) are not completely degraded in these early-replicating *B. bacteriovorus* cells. In *C. crescentus,* which also exhibits a biphasic life cycle, DnaA (DnaA_Cs_) undergoes cell cycle-controlled proteolysis mediated by the Lon protease (54, 55). The accumulation of DnaA_Cs_ in replication-active cells of *C. crescentus* corresponds to a low synthesis level of CtrA, which represses chromosome replication initiation (54, 55). Controlled proteolysis of DnaA and/or repression of chromosome replication by a CtrA-like protein could possibly occur in *B. bacteriovorus* during the attack phase. In this scenario, the level of this putative replication repressor might be too low to inhibit replication in newly released cells, and additionally, such cells could contain levels of replication proteins sufficient to restart chromosome replication. Unexpectedly, in few predators released in close proximity to new prey cells, the replisomes were not disassembled suggesting that in the higher population density of prey’s cells, the replication machinery might be “inherit”. We speculate that such predators can begin chromosome replication earlier in the next prey (see Fig. 4) than the cells that have to assemble replication machinery *de novo.* However, this hypothesis remains to be further elucidated.

In *B. bacteriovorus*, chromosome replication is not immediately followed by cell division; instead, a multinucleoid filament is formed. Such replication dynamics resembles that found in the vegetative and aerial mycelia of *Streptomyces* (56, 57). Moreover, after termination of replication, the multinucleoid filament (similar to the sporulating aerial hyphae of *Streptomyces*) undergoes synchronous septation (in up to 21 min after replication termination) to ensure that each nascent predatory cell receives a single copy of the chromosome. In contrast to the model organisms (*E. coli* and *B. subtilis*), *B. bacteriovorus* exhibits an extended B and D periods; the chromosome replication begins approximately 74 min after bdelloplast formation and is terminated before filament fragmentation inside bdelloplast.

To conclude, we herein show for the first time that the predatory cells of *B. bacteriovorus* exhibit an unusual spatiotemporal arrangement of chromosome replication dynamics that combine different features from Gram-negative and -positive bacteria. The chromosome replication of *B. bacteriovorus* initiates at the specific cell pole (the invasion one), as also seen in other asymmetrical bacteria, *C. crescentus* and *V*. *cholerae* (chromosome I). Interestingly, we observed ‘cell-to-cell’ variation in the replication dynamics. In a “rich” environment, i.e., in a dense prey cell population, the newly released, not fully matured predatory cells are able to quickly attack prey in the surrounding neighborhood and begin earlier the chromosome replications (see Tables 2 and S2). Surprisingly, in some DnaN-mNeonGreen tagged predatory cells, the replication machinery is not disassembled after septation. Moreover, in few cases, the replisome was present even during the attack phase what might allow predatory cell to restart chromosome replication shortly after bdelloplast formation. In larger prey cells that provide more nutrients, *B. bacteriovorus* grows as a long filament that exhibits high replication activity resulting in synthesis of more chromosomes (up to 12). On the other hand, in the case where *B. bacteriovorus* predation is abortive (e.g. due to the small size of prey, see Fig. 5), the predatory bacterium can complete its chromosome replication and consequently its cell cycle by encountering and invading another prey cell (Fig. 6). We speculate that heterogeneity in replication dynamics may reflect a relaxation of cell cycle checkpoints, possibly increasing the ability of predatory cells to adapt to different prey’s specific conditions-remembering that these predators replicate within a wide range of different prey genera. Thus, the population of *B. bacteriovorus* as other bacterial populations is not homogenous and some individuals can show unique features different from others.

## Materials and Methods

### DNA manipulations, bacterial strains and culture conditions

DNA manipulations in *E. coli* were carried out using standard protocols (58). Reagents and enzymes were supplied by Thermo Scientific and Sigma-Aldrich. Oligonucleotides were synthesized by Sigma-Aldrich. The plasmids used to construct *B. bacteriovorus* HD100 strain DnaN-mNeonGreen/PilZ-mCherry (see below) were propagated in *E. coli* DH5a, grown in LB broth or on LB agar plates (supplemented with 50 kanamycin μg/ml) and then transformed into *E. coli* S17-1. The latter were grown in liquid culture in YT medium (0.8% Bacto tryptone, 0.5% yeast extract, 0.5% NaCl, pH 7.5) with (S17-1:pZMR100) or without (S17-1) kanamycin (50 μg/ml), at 37°C with shaking (180 rpm). *B. bacteriovorus* was grown by predation on *E. coli* S17-1 or *E. coli* S17-1 pZMR100 (kanamycin-resistant strains) in Ca-HEPES buffer (25 mM HEPES, 2 mM calcium chloride, pH 7.6) as described in Lambert *et al.* (59). Details regarding the utilized strains, plasmids and oligonucleotides are listed in Table S1.

### Construction of *B. bacteriovorus* strain HD100 DnaN-mNeonGreen/PilZ-mCherry

We constructed *B. bacteriovorus* strain HD100 DnaN-mNeonGreen/PilZ-mCherry, in which the cytoplasm was labeled red by the PilZ fusion and the replisome labeled green by the DnaN fusion. We amplified the coding sequences of *dnaN* (primers pK18_dnaN(Gib)F and mNeon_dnaN(Gib)R) and *mNeonGreen* (primers dnaN_mNeon(Gib)F and pK18_mNeon(Gib)R) using chromosomal *B. bacteriovorus* HD100 and pAKF220 (plasmid kindly provided by Andrew K. Fenton), respectively, as the templates. Gibson assembly was used to clone the PCR products into *pK18mobsacB.* The obtained construct (pK18dnaN-*mNeonGreen*) was transformed into *E. coli* S17-1 and conjugated to *B. bacteriovorus* strain HD100 PilZ-mCherry as described previously (59). Single crossing-over of pK18dnaN-*mNeonGreen* into the *B. bacteriovorus* chromosome replaced the wild-type copy of *dnaN* with the DnaN-mNeonGreen fusion-encoding gene (Fig. S1A). From this, we obtained a *B. bacteriovorus* strain with the *dnaN-mNeonGreen* fusion under the control of the endogenous promoter and a second disrupted and a non-expressed copy of the *dnaN* gene. Proper construction of the DnaN-mNeonGreen/PilZ-mCherry strain was verified by PCR, sequencing and Western blotting.

### Time-Lapse Fluorescent Microscopy (TLFM)

Cells of *B. bacteriovorus* strain DnaN-mNeonGreen/PilZ-mCherry were prepared by predation on *E. coli* S17-1:pZMR100 in 50 ml Ca-HEPES buffer in the presence of 50 μg/ml kanamycin. The culture was spun down at 5500 rpm for 20 min at 30°C, resuspended in 5 ml of Ca-HEPES buffer and incubated at 30°C with 200 rpm shaking for 30 min. Agarose gel (1%) in Ca-HEPES buffer with or without novobiocin (final concentration, 200 μg/ml) was poured into a 35-mm glass-bottom μ-Dish (Ibidi) and allowed to solidify. The gel was removed from the dish, flipped over to bottom-up and coated with *E. coli* S17-1 overnight culture. Next, a few drops of *B. bacteriovorus* suspension was added on the *E. coli* coated surface and spread by inoculation loop. That way prepared agarose gel was back placed in a 35-mm glass-bottom μ-Dish bottom-down. Images were recorded every 1 or 5 min using a Delta Vision Elite inverted microscope equipped with a Olympus 100X/1.40 and a Cool SNAP HQ2-ICX285 camera. PilZ-mCherry was visualized with a mCherry (EX575/25; EM625/45) and ND (50%) filters with exposure time of 200 ms. DnaN-mNeonGreen was visualized with GFP (EX475/28; EM525/48) and ND (50%) filters with exposure time of 80 ms. Differential interference contrast (DIC) images were taken with ND (5%) filter and exposure time of 50 ms. The captured images were analyzed using the ImageJ Fiji suite (http://fiii.sc/Fiii).

TLFM experiments were done in three independent biological replicates.

## Supplementary materials

Supplementary material for this article may be found at…

## Acknowledgements

This study was supported by National Science Centre, Preludium grant 2016/23/N/NZ1/02965 to LM; ERASMUS Traineeship (LM started initial microscopy an genetic manipulation in the laboratory of RES, 2016); BBSRC UK grant BB/M010325/1 to RT and CL; Leverhulme Trust UK grant RPG-2014-241 to RL. We thank Andrew K. Fenton for providing the pAKF220 plasmid and Luke Ray (RES lab) for helpful discussions.

