## Supplemental Materials for "Dynamics of chromosome replication and its relationship to predatory attack lifestyles in *Bdellovibrio bacteriovorus*"

**Supplementary Materials**

**Supplementary Materials and Methods**

**Prey survival assay of *B. bacteriovorus* strains**

Cells of *B. bacteriovorus* strains DnaN-mNeonGreen/PilZ-mCherry and wild-type were prepared by predation on *E. coli* S17-1 in 50 ml Ca-HEPES buffer. The cultures were filtered through 0.45 µm filters, spun down at 6000 rpm for 20 min at 30°C and resuspended in 510 µl of Ca-HEPES buffer giving the final concentration of approximately 1x$10^9$ pfu/ml (plaque-forming unit/ml). *E. coli* S17-1 overnight culture was spun down at 6000 rpm for 10 min at 20°C, washed with Ca-HEPES buffer and diluted to $OD_{600}$=1.0 with Ca-HEPES buffer giving the final concentration of approximately 1x$10^9$ cfu/ml. The assay was set up by adding 500 µl of *B. bacteriovorus* cells (DnaN/PilZ or wild-type) or Ca-HEPES buffer (control) to 5 ml of *E. coli* S17-1 (OD=1.0). Cultures were incubated at 30°C with 200 rpm shaking. To enumerate predatory cells concentration taken to predation assay dilutions of *B. bacteriovorus* cells were plated on overlay agar plates. Plates were incubated at 30°C for 5 days. To enumerate colony forming units (cfu) of prey cells, samples were taken at 0h, 3h, 6h, and 24h and plated in triplicate by Misra and Miles technique on YT agar plates at serial dilutions. Plates were incubated overnight at 37°C and then, visible *E. coli* colonies were enumerated. Pray survival assay was done in two independent biological replicates.

**Predatory kill curves of *B. bacteriovorus* strains**

*B. bacteriovorus* strains (DnaN-mNeonGreen/PilZ-mCherry and wild-type) were prepared as described above. *E. coli* S17-1 overnight culture was spun down at 6000 rpm for 10 min at 20°C, washed with Ca-HEPES buffer and diluted to $OD_{600}$ = 1.0 with Ca-HEPES buffer. 20 µl of filtrated *B. bacteriovorus* cells were added to 280 µl of *E. coli* suspension. Lysis curves were analyzed using Bioscreen C (Automated Growth Curves Analysis System, Growth Curves USA) by measuring the decrease of optical density ($OD_{600}$) at 30°C in 20-minute intervals for 27 hours. Experiments were done in three independent biological replicates.


**Legends to Supplementary Figures**

**Figure S1**

**Characteristics of the *B. bacteriovorus* strain, DnaN-mNeonGreen/PilZ-mCherry**

**(A)** Cartoon of the single crossover integration of pK18*dnaN-mNeonGreen* into the *B. bacteriovorus* chromosome. Black arrows indicate gene promotors. **(B)** PilZ-mCherry and

DnaN-mNeonGreen in *B. bacteriovorus* attack-phase cells, as assessed under epifluorescence microscopy. The following are shown (beginning on the left): differential interference contrast (DIC, indicated as "Phase") image; red fluorescence image; green fluorescence image; and merged phase and fluorescence images. **(C)** Effect of novobiocin on the presence of replisome foci. Novobiocin (200 µg/ml, final concentration) was added to 1% agarose gel in Ca-HEPES buffer, and the predation of *B. bacteriovorus* DnaN-mNeonGreen/PilZ-mCherry on *E. coli* S17-1 cells was observed using time-lapse fluorescence microscopy. Left, red fluorescence image; middle, green fluorescence image; right, merged phase and fluorescence images. Scale bar = 1 µm.

**Figure S2**

**Predatory kill curve and predation efficiency of B. bacteriovorus wild-type and DnaN-mNeonGreen/PilZ-mCherry**

**(A)** Predation kill curve of *B. bacteriovorus* strains

**(B)** Cfu/ml of *E. coli* cells after incubation with examining *B. bacteriovorus* strains. In red pfu/ml of *B. bacteriovorus* cells.

**Figure S3**

***B. bacteriovorus* entry and invasive pole localization in *E. coli* periplasm**

Attachment of *B. bacteriovorus* to *E. coli* host cell (**A**). Predatory entry into prey's periplasm in relation to invasive pole (**B-D**). Initiation of chromosome replication at invasive pole (**E**).

Red - PilZ-mCherry labelled cytoplasm of *Bdellovibrio* and green - DnaN-mNeonGreen of *Bdellovibrio.* Photos represent merged DIC and fluorescence (red and green) images. Pictures were taken every 1 min. Scale bar = 1 µm.

**Figure S4**

***B. bacteriovorus* growth and chromosome replication in an abnormally elongated *E. coli* cell**

Attachment of *B. bacteriovorus* to an elongated E. coli cell (**A**). Bdelloplast formation contracting and rounding the large *E.coli* cell, time = 0 min (**B**). Initiation (**C**) and continuation of chromosome replication (**D-E**). Progeny cells (**F**). The yellow arrow highlights the single predatory cell that invades the *E. coli* cell. The white arrows indicate the positioning of replisomes in the *B. bacteriovorus* filament.

Red - PilZ-mCherry labelled cytoplasm of *Bdellovibrio* and green - DnaN-mNeonGreen of *Bdellovibrio.* Photos represent merged DIC and fluorescence (red and green) images. The *B. bacteriovorus* and *E. coli* cells are marked by yellow and white dotted lines, respectively as determined by careful analysis of DIC images. Scale bar = 1 µm.

**Fig S5**

**Amino acid sequence alignments of DnaN protein from *E. coli* K-12 and *B. bacteriovorus* HD100**

Relatively high (55%) homology between *E. coli* K-12 and *B. bacteriovorus* HD100 DnaN proteins. Residues identical (or missed) for the *E. coli* and *B. bacteriovorus* DnaN sequences are printed in black background; residues not identical but at least similar are printed in gray background.

**Figure S6**

**Amino acid sequence alignments of DnaE protein (catalytic alpha subunit of DNA polymerase III) from *E. coli* K-12 and *B. bacteriovorus* HD100**

Relatively high (47%) homology between *E. coli* K-12 and *B. bacteriovorus* HD100 DnaE proteins. Residues identical (or missed) for the *E. coli* and *B. bacteriovorus* DnaN sequences are printed in black background; residues not identical but at least similar are printed in gray background. Amino acid residues crucial for *E. coli* DnaE enzyme activity are indicated in red (1).

**Legends to Supplementary Movies**

**Movie S1** Time-laps imaging of *B. bacteriovorus* entry into *E. coli* periplasm in relation to invasion pole and subcellular localization of DnaN-mNeonGreen (green) in strain HD100 DnaN-mNeonGreen/PilZ-mCherry. Predatory cell indicated by PilZ-mCherry (red). Differential interference contrast (grey) signals were taken every 1 min.

**Movie S2** Time-lapse imaging of replisomes in *B. bacteriovorus*. Subcellular localization of DnaN-mNeonGreen (green) in strain HD100 DnaN-mNeonGreen/PilZ-mCherry. Predatory cell indicated by PilZ-mCherry (red). Differential interference contrast (grey) signals were taken every 5 min.

**Movie S3** Time-laps imaging of replisomes in *B. bacteriovorus* growing in abnormally elongating *E. coli* cell. Subcellular localization of DnaN-mNeonGreen (green) in strain HD100 DnaN-mNeonGreen/PilZ-mCherry. Predatory cell indicated by PilZ-mCherry (red). Differential interference contrast (grey) signals were taken every 5 min.

**Movie S4** Time-lapse imaging of newly-released *B. bacteriovorus* cell with visible fluorescence focus at the invasive pole during attack phase. Subcellular localization of DnaN-mNeonGreen (green) in strain HD100 DnaN-mNeonGreen/PilZ-mCherry. Predatory cell indicated by PilZ-mCherry (red). Differential interference contrast (grey) signals were taken every 5 min.

**Movie S5** Time-laps imaging of *B. bacteriovorus* chromosome growth and replication in two independent host cells. Subcellular localization of DnaN-mNeonGreen (green) in strain HD100 DnaN-mNeonGreen/PilZ-mCherry. Predatory cell indicated by PilZ-mCherry (red). Differential interference contrast (grey) signals were taken every 5 min.
