## Supplemental Tables for "Dynamics of chromosome replication and its relationship to predatory attack lifestyles in *Bdellovibrio bacteriovorus*"

**Table S1** Bacterial strains, primers and plasmids

**Bacterial strains**

| Strain | Description | Reference/source |
|---|---|---|
| ***E. coli*** | | |
| S17-1 | *thi pro hsdR⁻hsdM⁺recA*; integrated plasmid RP4-Tc::Mu-Kn::Tn7 used as donor for conjugation of plasmids into *Bdellovibrio* | (2) |
| S17-1 pZMR100 | S17-1 strain containing pZMR100 plasmid to confer $Kana^R$; used as $Kana^R$ prey for *Bdellovibrio* | (3) |
| ***B. bacteriovorus*** | | |
| HD100*Bd0064-mCherry* | Merodiploid HD100 with wild-type Bd0064 fused with mCherry. | (4) |
| HD100*Bd0064-mCherry/Bd0002-mNeon* | HD100*Bd0064-mCherry* strain carrying integrated plasmid pK18_mNeon_dnaN at the *dnaN* (Bd0002) locus | This work |

**Primers**

| Name | 5' – 3' sequence |
|---|---|
| pK18_dnaN(Gib)F | CGTTGTAAAACGACGGCCAGTGCCAATGAAATTAGAGATTGATAAGCG |
| mNeon_dnaN(Gib)R | CTTTCGAAACCATGATTCTCATTGGCATCAC |
| dnaN_mNeon(Gib)F | GCCAATGAGAATCATGGTTTCGAAAGGAGAG |
| pK18_mNeon(Gib)R | GGAAACAGCTATGACCATGATTACGTCACTTATAGAGTTCATCCATACC |

**Plasmids**

| Name | Feature | Reference/source |
|---|---|---|
| pAKF220 | Plasmid carrying *mNeonGreen* sequence; $Amp^R$ | Andrew K. Fenton |
| pK18*mobsacB* | Suicide vector used for conjugation and recombination into *Bdellovibrio* genome; $Kana^R$ | (5) |
| pK18_dnaN_mNeon | Derivative of pK18*mobsacB* containing C-terminal fused *mNeonGreen* to *bd0002 (dnaN)*; $Kana^R$ | This work |

**Table 2** Comparison of average time of replisomes appearing in free-living and newly-released *B. bacteriovorus* cells

| | Replisome appearance (mean±SD) [min] | | | | Time intervals (mean±SD) [min] | | | Total replication time (mean±SD) [min] |
|---|---|---|---|---|---|---|---|---|
| | **I** | **II** | **III** | **IV** | **I → II** | **II → III** | **III → IV** | |
| **Free-living cells** | 74±26 | 133±32 | 164±33 | 177±29 | 59±20 | 32±18 | 27±15 | 144±26 |
| **Newly-released cells** | *** 23±11 | 77±21 | 103±15 | 128±17 | 53±16 | 26±15 | 26±13 | 140±20 |

*** p-value < 0.001
