## Supplementary figures and images for "Dynamics of chromosome replication and its relationship to predatory attack lifestyles in *Bdellovibrio bacteriovorus*"

### Supplemental Figure 1

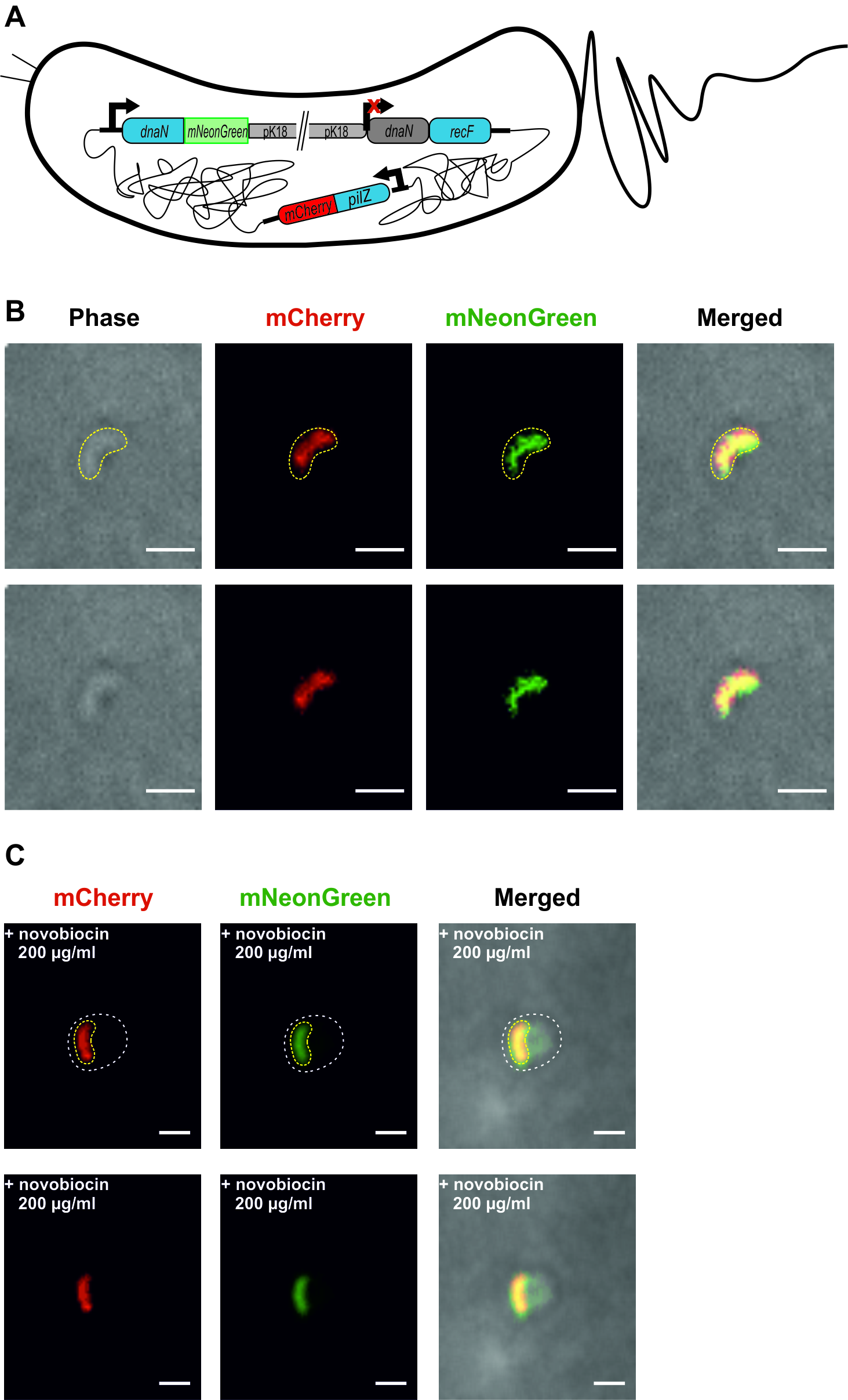

### Supplemental Figure 2

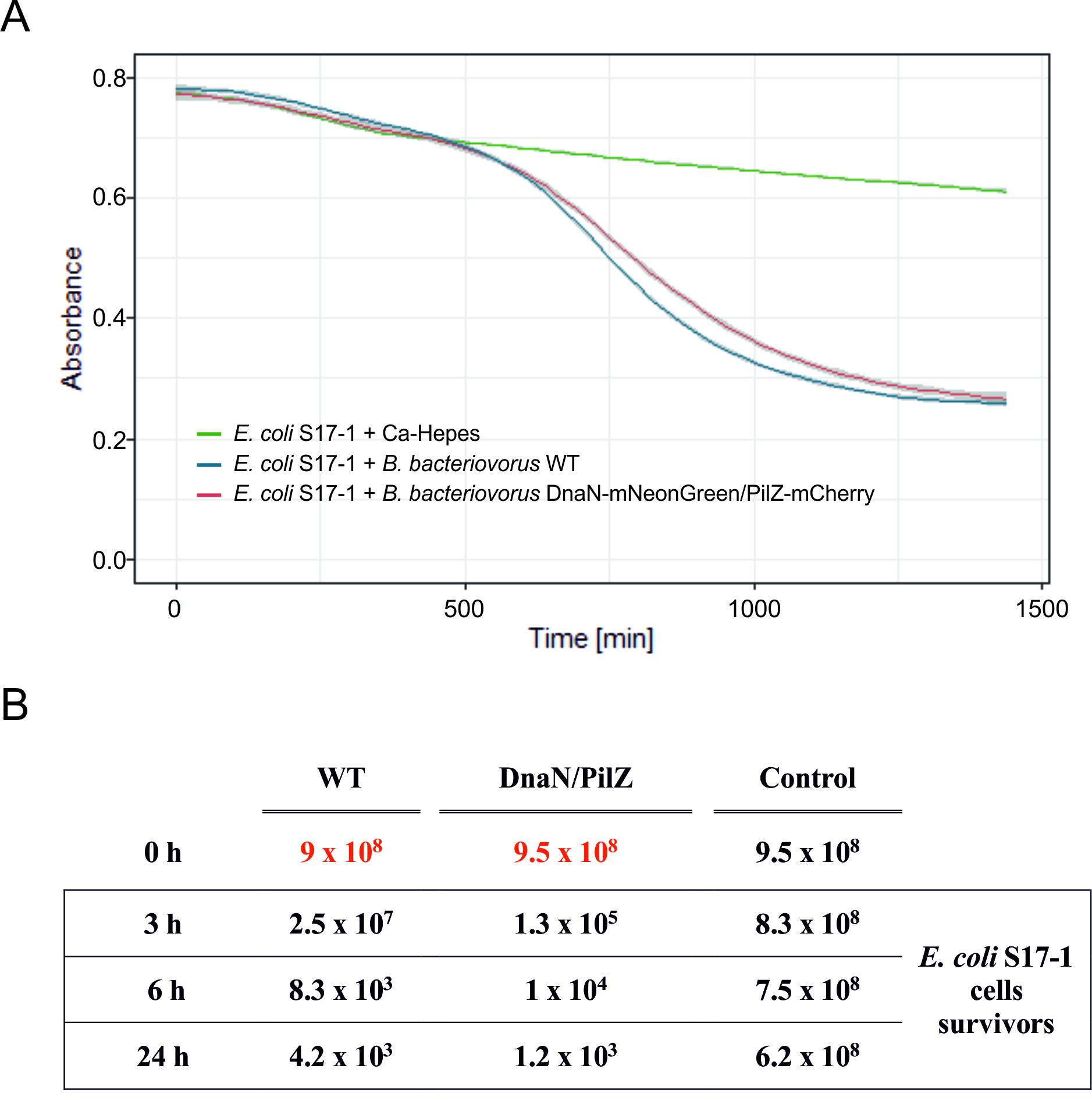

### Supplemental Figure 3

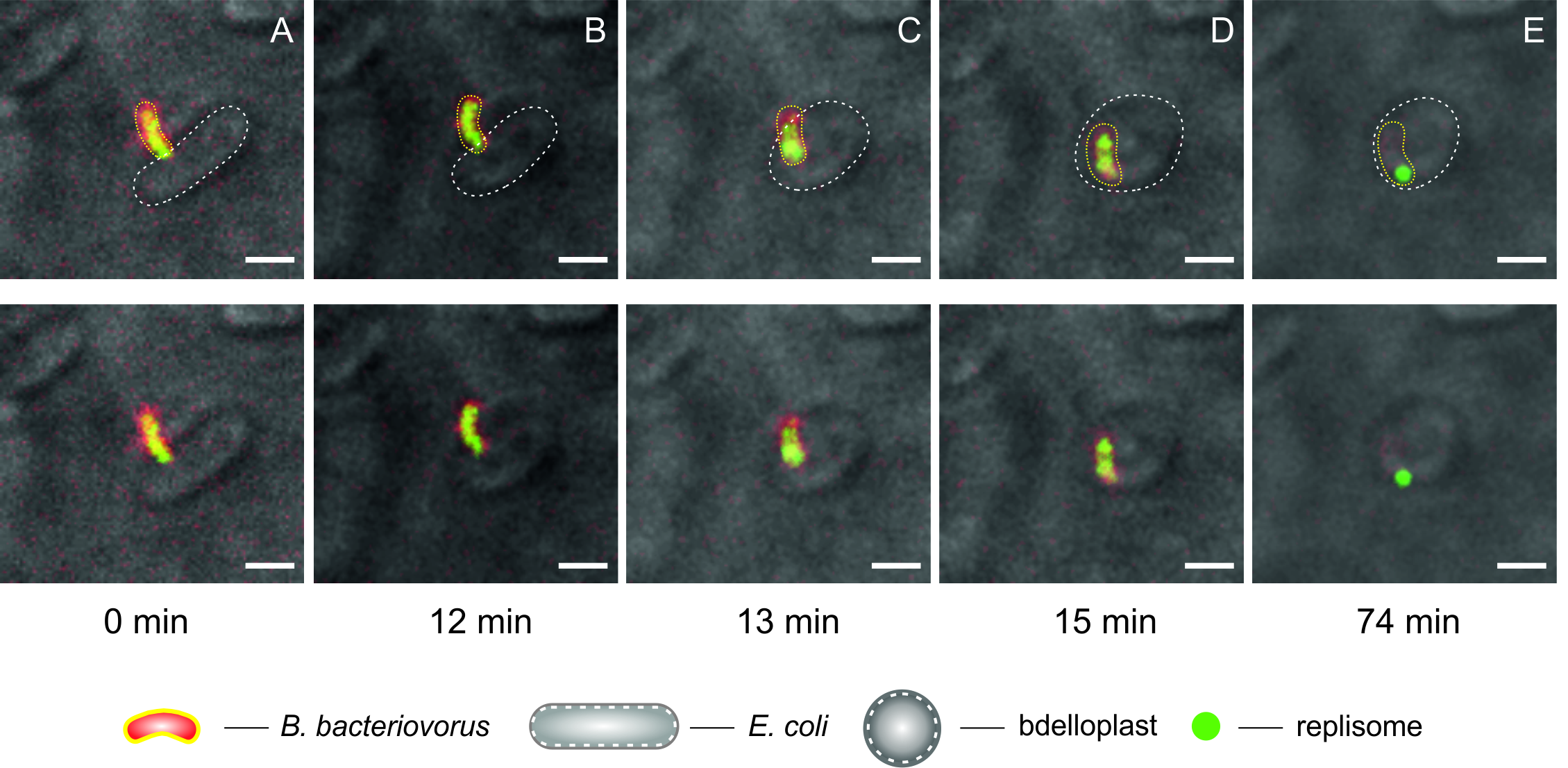

### Supplemental Figure 4

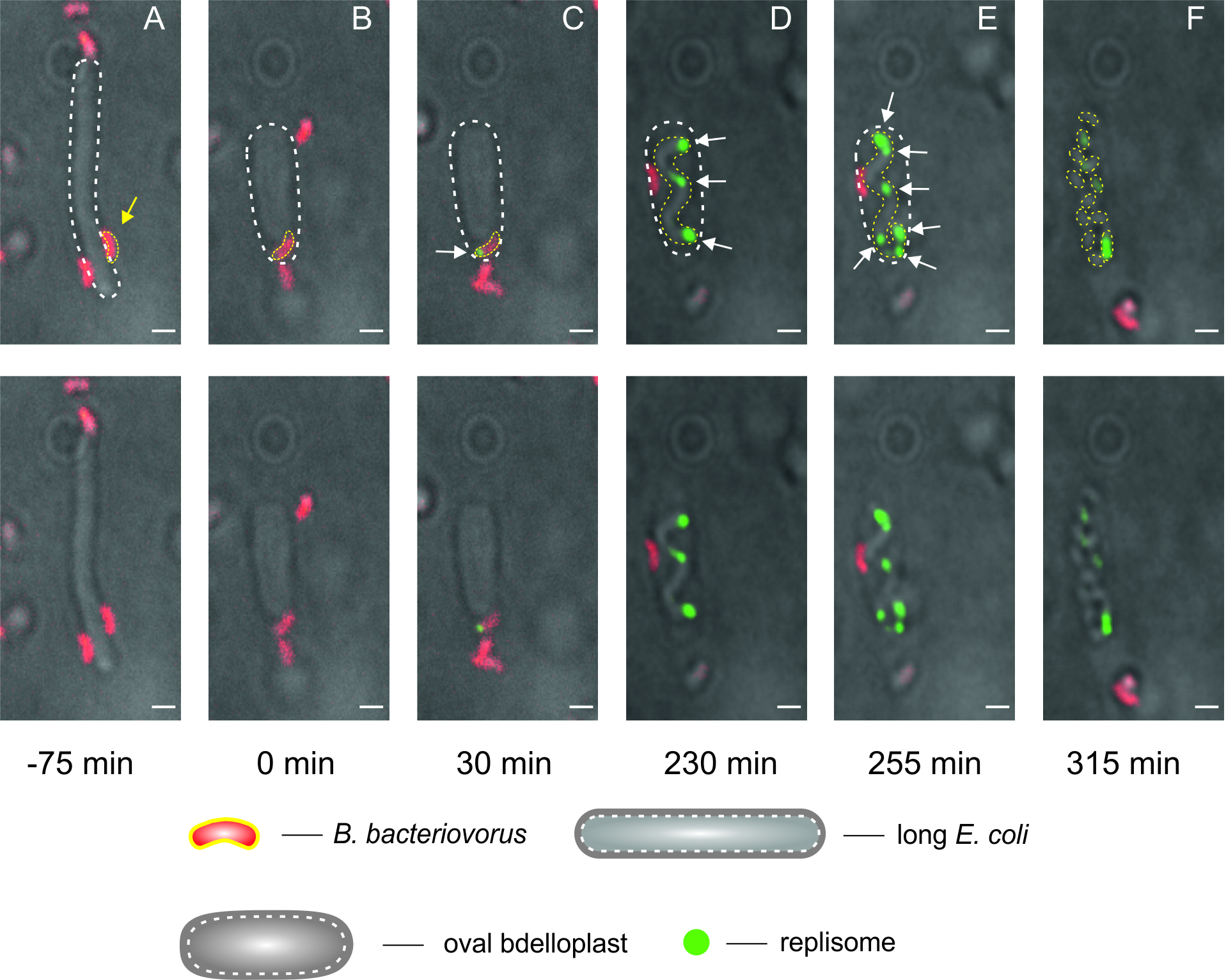

### Supplemental Figure 5

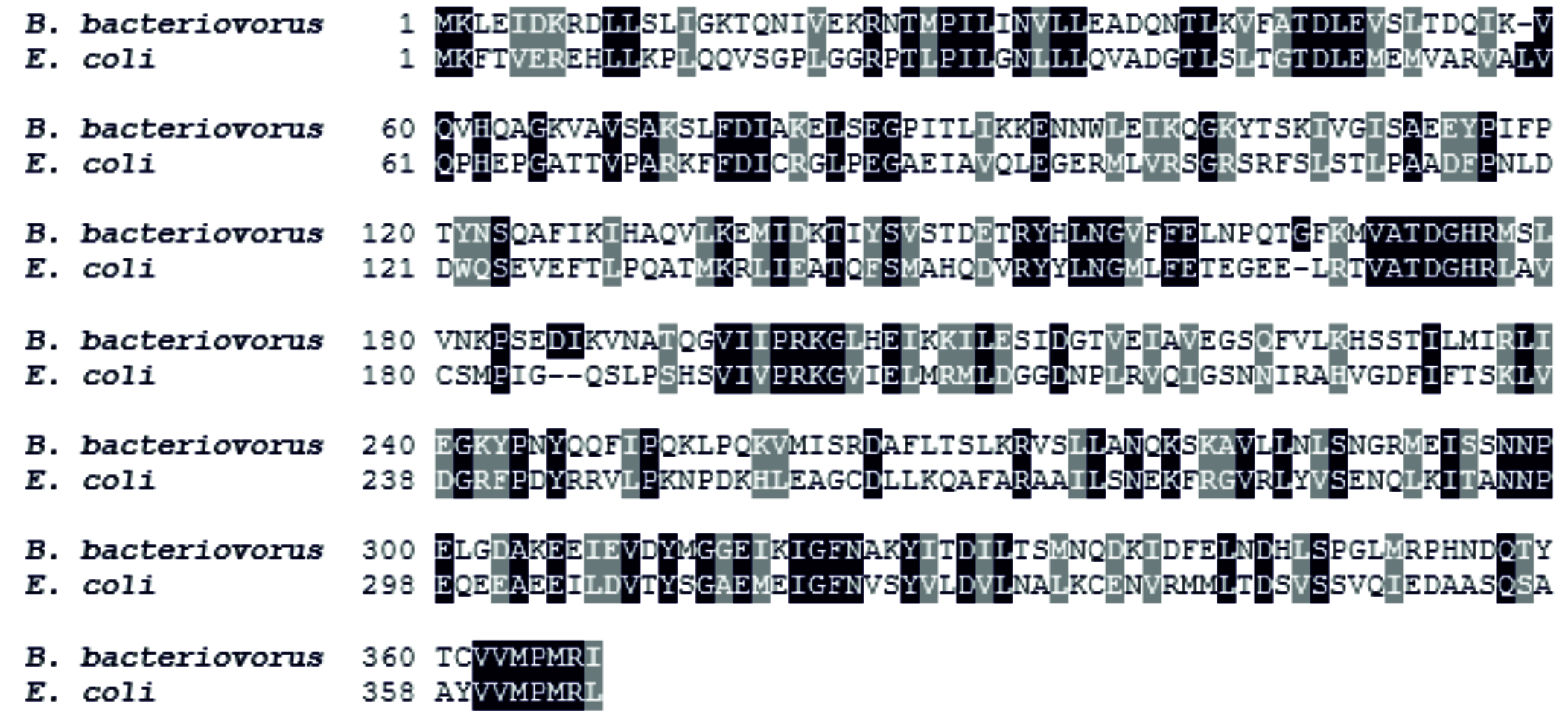

### Supplemental Figure 6

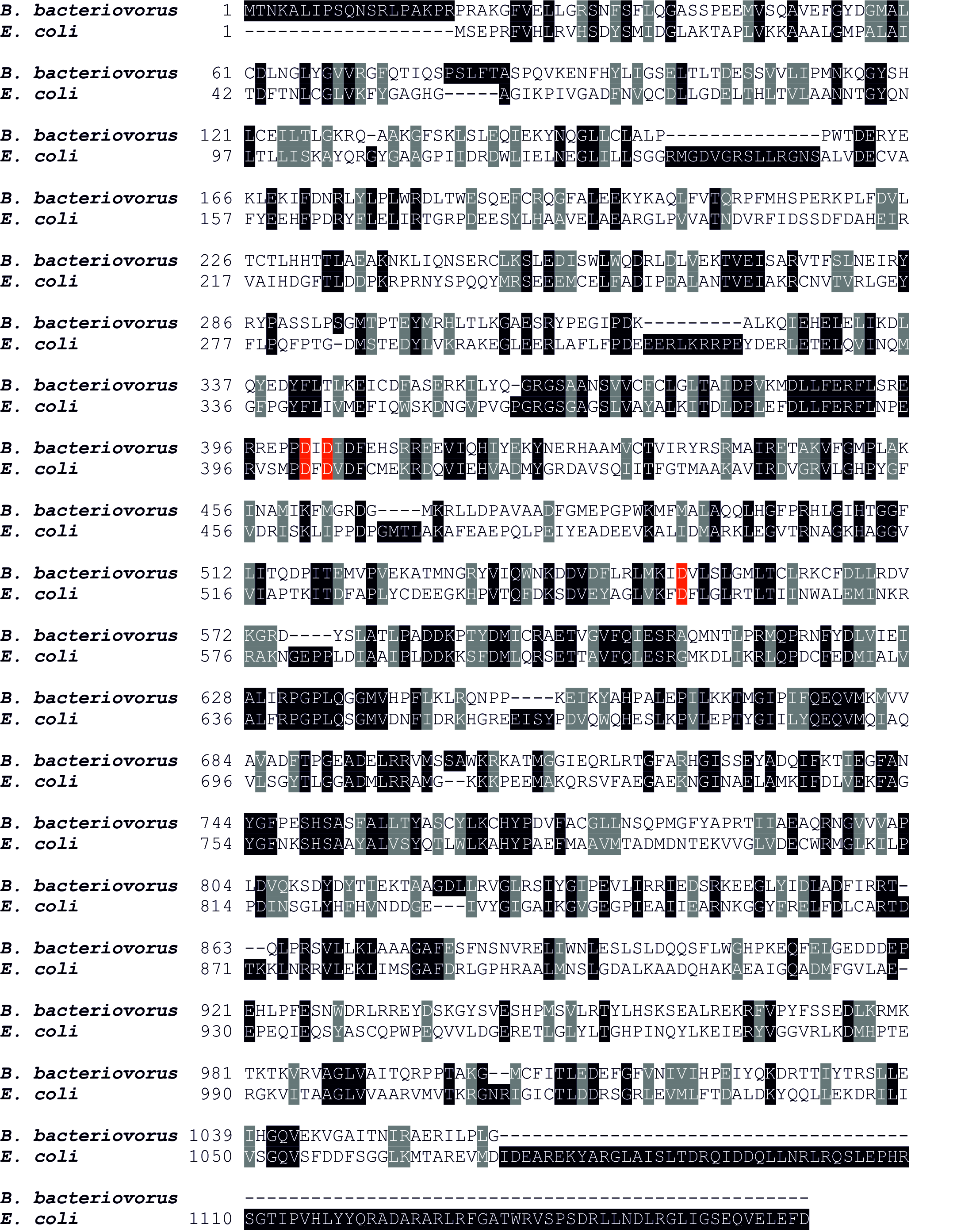
